# The association of plasma lipids with white blood cell counts: Results from the Multi-Ethnic Study of Atherosclerosis

**DOI:** 10.1101/488023

**Authors:** Yong Chang Lai, Kevin J. Woollard, Robyn L. McClelland, Matthew A. Allison, Kerry-Anne Rye, Kwok Leung Ong, Blake J. Cochran

## Abstract

**Background:** Previous studies have demonstrated that elevated cholesterol results in increased white blood cell counts in mouse models. However, there is insufficient evidence to support this in humans.

**Objective:** To investigate the relationship of plasma lipids with white blood cell counts (basophils, eosinophils, monocytes, neutrophils and lymphocytes) in the Multi-Ethnic Study of Atherosclerosis (MESA).

**Methods:** The analysis included 2873 MESA participants with a complete white blood count and differential analysis. The cross-sectional association of total cholesterol, LDL cholesterol, HDL cholesterol and triglyceride levels with different white blood cell counts was analyzed by multivariable linear regression.

**Results:** After adjusting for sociodemographic and confounding factors including red blood cell counts, platelet counts, use of lipid-lowering medication, CVD risk factors and other lipid measures, and multiple testing correction, a 1-SD increment in total cholesterol and LDL cholesterol was associated with 2.8% and 2.3% lower total white blood cell counts, 3.7% and 3.0% lower monocyte counts, and 3.4% and 2.7% lower neutrophil counts (all *p*<0.01). The same increment in ln-transformed triglyceride levels was associated with 2.3% higher total white blood cell counts and 4.5% higher lymphocyte counts (both *p*<0.001). Similar results were obtained after excluding participants taking lipid-lowering medication. A 1-SD increment increase in HDL cholesterol was associated with a 1.5% lower white blood cell count (*p*=0.018), but was not significantly associated with changes in any individual cell type.

**Conclusion:** Whilst significant associations were observed between plasma lipid levels and white blood cell populations, the heterogenous and modest nature of these relationships make it hard to support the hypothesis that lipids are in the causal pathway for leukogenesis in humans.

## Introduction

Leukocytes play important roles in immune regulation and tissue homeostasis^1^. Despite studies in animal models demonstrating a link between dyslipidaemia and increased leucocytosis, the impact of high plasma lipid levels on white blood cell counts in humans is poorly understood. With respect to animal models, Murphy, et al.^2^ showed that accumulation of cholesterol in hematopoietic stem progenitor cells in apoE^-/-^ mice increased responsiveness to IL-3 and granulocyte-macrophage colony-stimulating factor. This resulted in progenitor cell proliferation, leading to neutrophilia and monocytosis. Hypercholesterolaemia can also increase circulating levels of C-X-C motif ligand 1 (CXCL1), and increase expression of C-X-C chemokine receptor type 2 (CXCR2) in neutrophils, thereby inducing neutrophil release from bone marrow^3^.

A previous study of children with familial hypercholesterolaemia reported no relationship between cholesterol levels and neutrophil counts, but did identify a significant relationship between elevated HDL cholesterol levels and decreased monocyte numbers^4^. A recent study by Andersen et al.^5^ using data from the United States National Health and Nutritional Examination Survey (NHANES) found significant positive associations between triglyceride levels and lymphocyte, neutrophil, monocyte and basophil counts as well as sex-dependent inverse relationships between HDL cholesterol levels and leukocyte counts. Increased triglyceride (as remnant lipoproteins) has also been associated with increased total white blood cell, monocyte, lymphocyte and neutrophil counts^6^, and increased LDL cholesterol levels are associated with increased lymphocyte number^7, 8^. A modest positive association between non-HDL cholesterol levels and platelet counts has also been reported^9^.

It may also be important to consider the impact of dyslipidaemia on cellular function beyond simply influencing cell number. Changes in cholesterol homeostasis have been associated with alterations in monocyte subsets in humans^10^, and increased cellular cholesterol levels has been demonstrated to increase proliferation^11, 12^ and inflammatory responses^13^ in T lymphocytes.

As both increased lipids and the number of myeloid cells can have an impact on cardiovascular disease (CVD) risk, it is important to determine and understand the interactions between these factors. We therefore investigated the relationship of different lipid measures with different types of white blood cell counts, and whether there were any sex or racial/ethnic differences in the relationship, using data from the Multi-Ethnic Study of Atherosclerosis (MESA)^14^.

## Materials and Methods

### Study participants

The Multi-Ethnic Study of Atherosclerosis (MESA) was designed to investigate the prevalence, risk factors and progression of subclinical cardiovascular disease (CVD) in a multi-ethnic cohort. Details of the study have been described previously^14^. Briefly, the study recruited 6814 men and women aged 45-84 years from four different ethnic groups, non-Hispanic whites, African American, Hispanic American and Asian (predominantly of Chinese descent), between July 2000 and August 2002. These participants were recruited from six different US communities (Baltimore, MD; Chicago, IL; Forsyth County, NC; Los Angeles County, CA; New York, NY, and St. Paul, MN) and were all free of clinically apparent CVD at baseline (exam 1). The study was approved by the institutional review boards at all participating centres and undertaken in accordance with the Declaration of Helsinki and Good Clinical Practice Guidelines. Informed written consent was obtained from all participants. This study was approved by the UNSW Human Research Advisory Panel (HC180331).

The participants underwent four follow up assessments (exams 2-5) over a period of 10 years. Total white blood cell and subfraction counts were measured only at exam 5 (2010-2012). Of the 6814 participants at baseline exam 1, 4716 participants attended exam 5, 2890 of whom had their white blood cell counts measured. After excluding 17 participants with missing data in one or more measures of plasma lipids (total cholesterol, HDL cholesterol, LDL cholesterol and triglycerides) at exam 5, a total of 2873 participants were included in the analysis.

### Lipid and white blood cell measurements

Venous blood samples were collected in the morning after a 12-hour overnight fast by certified technicians using standardised venepuncture procedures. Total cholesterol, HDL cholesterol, LDL cholesterol and triglycerides, as well as basophil, eosinophil, lymphocyte, monocyte, neutrophil and total white blood cell counts, were measured as described previously^15, 16^. Apolipoprotein E *(APOE)* genotype was estimated from single nucleotide polymorphisms rs429358 and rs7412 as described previously^17^. Serum glucose was measured by rate reflectance spectrophotometry using thin film adaptation of the glucose oxidase method. Diabetes was defined as fasting blood glucose ≥7.0 mmol/L (126 mg/dL) or use of hypoglycemic medication^18^. Serum creatinine was measured via rate reflectance spectrophotometry and calibrated to the standardised creatinine measurements obtained at the Cleveland Clinic Research Laboratory (Cleveland, OH). Estimated glomerular filtration rate (eGFR) was calculated from serum creatinine using the Chronic Kidney Disease Epidemiology Collaboration (CKD-EPI) Creatinine Equation^19^.

### Other covariates of interest

At MESA exam 5, information about demographics, socioeconomic status, medication, dietary and alcohol intake, smoking status, pack-years of smoking, physical activity and medical history were collected via self-reported questionnaire. History of CVD was defined as the presence of an adjudicated hard CVD event, including myocardial infarction, resuscitated cardiac arrest, and stroke prior to, or at the time of, exam 5. History of coronary revascularization was defined as having percutaneous coronary intervention or coronary artery bypass grafting prior to, or at the time of, exam 5. Details of the CVD and coronary revascularization event ascertainment has been described previously^20, 21^.

The height and weight of participants was measured to the nearest 0.1 cm and 0.5 kg, respectively, with participants wearing light clothing and no shoes. Body mass index (BMI) was calculated by dividing the height in metres squared by the weight in kilograms. Waist and hip circumference were measured in centimetres using a steel measuring tape. Resting blood pressure was measured in the right arm after 5 min in the seated position using an automated oscillometric method (Dinamap) and appropriate cuff size. Readings were taken three times, with the last two values averaged to obtain the blood pressure reading that was used for analysis. Hypertension was defined as blood pressure ≥140/90 mmHg or taking antihypertensive medication^22^. Inflammation and infectious-related conditions were defined as self-reported history of fever, cold/flu, urinary infection, seasonal allergy, bronchitis, sinus infection, pneumonia, tooth infection or impaired liver function over the past two weeks.

### Statistical analysis

SPSS 24 (IBM, Armonk, NY) and STATA 14.0 (StataCorp, College Station, TX) were used for statistical analysis. For variables with a skewed distribution (including triglycerides and different types of white blood cell counts), the data was presented as median (interquartile range) and subjected to natural logarithmic (ln) transformation before analysis to improve normality. For cell counts with a zero value, such as basophils and eosinophils, a constant of 1 was added to the values before ln-transformation.

Multivariable linear regression analysis was performed using robust standard error estimation with each lipid measure (total cholesterol, HDL cholesterol, LDL cholesterol and triglycerides) as independent predictor variables, and basophil, eosinophil, lymphocyte, monocyte, neutrophil and total white blood cell counts as the dependent variables. The regression coefficient (B) was estimated as the ln-transformed cell counts per one SD unit increase in lipid levels. In model 1, data were adjusted for demographic factors including age, sex, and race/ethnicity. In model 2, data were further adjusted for cardiovascular risk factors including BMI, family income, smoking, pack-years of smoking, current alcohol use, physical activity, hypertension, diabetes, use of any lipid-lowering medication, eGFR, *APOE* genotype, history of CVD and history of coronary revascularization. In model 3, data were further adjusted for total cholesterol, HDL cholesterol, LDL cholesterol, and triglycerides, where appropriate. In model 4, data were further adjusted for other blood cell count related variables, including red blood cell and platelet counts. In all of the analyses, no multi-collinearity issue was detected (all variance inflation factors <3.0). Multiple testing correction was performed using false discovery rate (FDR) with the study-wide FDR at 0.05.

Analysis of diagnostic residual plots suggested the normality of the model residuals was improved after ln-transformation of total white blood cell, monocyte, neutrophil and lymphocyte counts, but not for basophil and eosinophil counts. For these two blood cell types, ln-transformation was used in the main analysis to be consistent with other blood cell types and a previous MESA study^16^. Therefore, a sensitivity analysis was performed without ln-transformation for basophil and eosinophil counts. In another sensitivity analysis, the above analyses were repeated after further excluding 1158 participants taking lipid-lowering medication.

In a separate analysis, we assessed the non-linear relationship of lipid measures with different types of white blood cell counts using the STATA command “mvrs” which uses regression splines to model potentially nonlinear relationships. We allowed the relationship for the lipid exposure of interest to be nonlinear while assuming that the relationships of other confounding variables with different white blood cell counts were linear. In two situations where nonlinearity was present the approximate knot positions were used to fit linear regression analyses within strata defined by these thresholds.

Interactions with sex and race/ethnicity was estimated by including the interaction term in the regression model in the full sample after adjustment for the main effects of the covariates. However, as the number of Chinese Americans was too small to make inferences about interaction, Chinese Americans were excluded from analysis of race/ethnicity interaction. Subgroup analysis was performed if there was a significant interaction. A two-tailed *p*<0.05 was considered to be statistically significant.

## Results

### Baseline Characteristics

Table 1 shows baseline characteristics of the 2873 participants. The cohort consisted of 42.8% non-Hispanic White Americans, 0.9% Chinese-Americans, 27.9% African-American and 28.3% Hispanic-Americans. The mean age was 69.5 years and 52.9% were women. The mean total cholesterol, HDL cholesterol and LDL cholesterol levels were 180.8, 55.0 and 104.4 mg/dL respectively, whereas the median triglyceride levels were 93 mg/dL. The median total white blood cell counts levels were 5.8×10^3^/μL. As shown in Supplementary Table 1, participants included in this analysis were younger, less likely to be Chinese Americans, and more likely to be non-Hispanic White or Hispanic Americans, compared to those participants not included in this study. Supplementary Figure 1 also shows the distribution of different blood counts with and without ln-transformation.

**Table 1.**
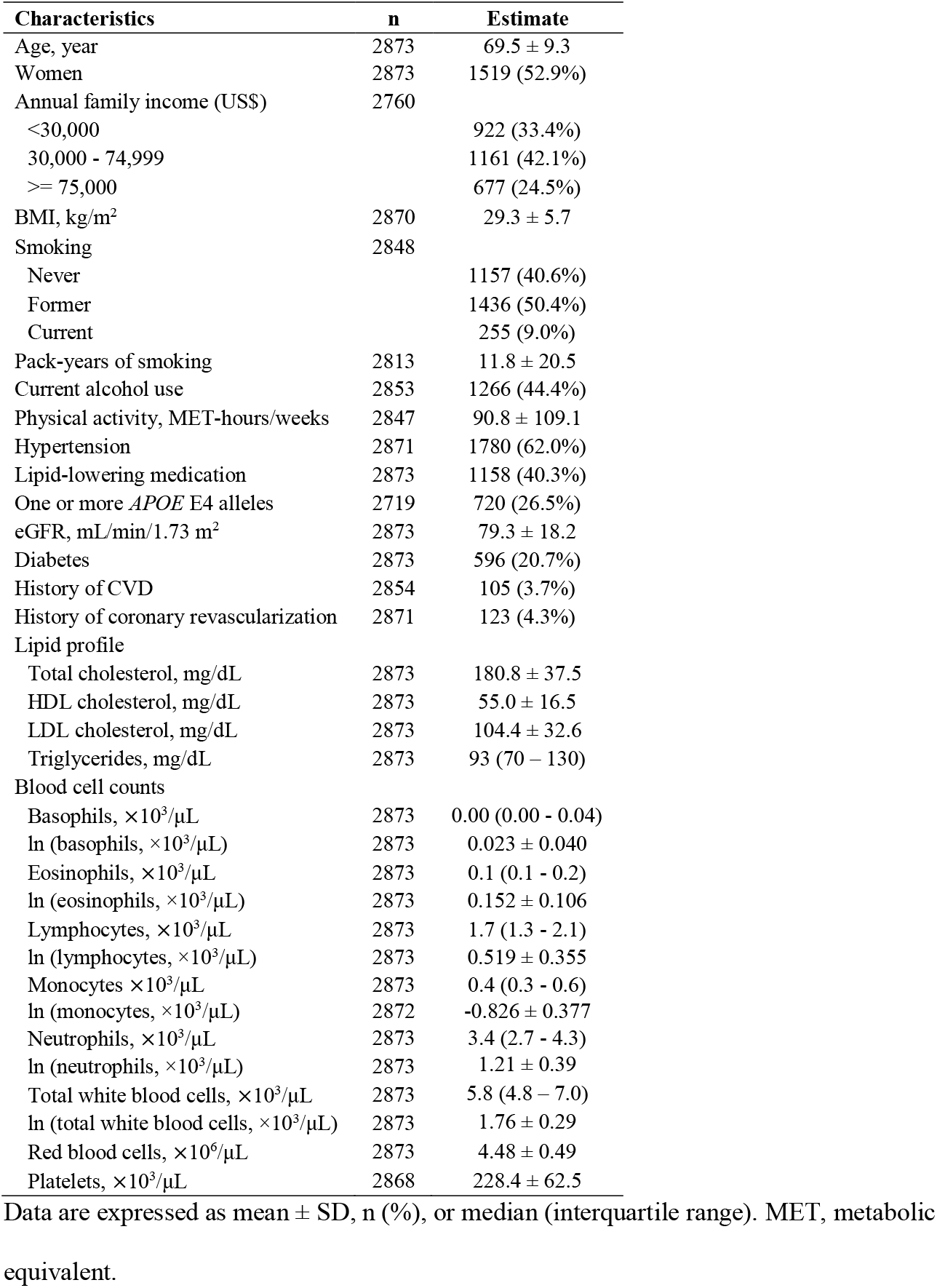
Baseline characteristics of participants

| Characteristics | n | Estimate |
| --- | --- | --- |
| Age, year | 2873 | 69.5 ± 9.3 |
| Women | 2873 | 1519 (52.9%) |
| Annual family income (US\$) | 2760 | |
| <30,000 |  | 922 (33.4%) |
| 30,000 - 74,999 |  | 1161 (42.1%) |
| >= 75,000 |  | 677 (24.5%) |
| BMI, kg/m<sup>2</sup> | 2870 | 29.3 ± 5.7 |
| Smoking | 2848 |  |
| Never |  | 1157 (40.6%) |
| Former |  | 1436 (50.4%) |
| Current |  | 255 (9.0%) |
| Pack-years of smoking | 2813 | 11.8 ± 20.5 |
| Current alcohol use | 2853 | 1266 (44.4%) |
| Physical activity, MET-hours/weeks | 2847 | 90.8 ± 109.1 |
| Hypertension | 2871 | 1780 (62.0%) |
| Lipid-lowering medication | 2873 | 1158 (40.3%) |
| One or more *APOE* E4 alleles | 2719 | 720 (26.5%) |
| eGFR, mL/min/1.73 m<sup>2</sup> | 2873 | 79.3 ± 18.2 |
| Diabetes | 2873 | 596 (20.7%) |
| History of CVD | 2854 | 105 (3.7%) |
| History of coronary revascularization | 2871 | 123 (4.3%) |
| Lipid profile |  |  |
| Total cholesterol, mg/dL | 2873 | 180.8 ± 37.5 |
| HDL cholesterol, mg/dL | 2873 | 55.0 ± 16.5 |
| LDL cholesterol, mg/dL | 2873 | 104.4 ± 32.6 |
| Triglycerides, mg/dL | 2873 | 93 (70 – 130) |
| Blood cell counts |  |  |
| Basophils, ×10<sup>3</sup>/μL | 2873 | 0.00 (0.00 - 0.04) |
| ln (basophils, ×10<sup>3</sup>/μL) | 2873 | 0.023 ± 0.040 |
| Eosinophils, ×10<sup>3</sup>/μL | 2873 | 0.1 (0.1 - 0.2) |
| ln (eosinophils, ×10<sup>3</sup>/μL) | 2873 | 0.152 ± 0.106 |
| Lymphocytes, ×10<sup>3</sup>/μL | 2873 | 1.7 (1.3 - 2.1) |
| ln (lymphocytes, ×10<sup>3</sup>/μL) | 2873 | 0.519 ± 0.355 |
| Monocytes ×10<sup>3</sup>/μL | 2873 | 0.4 (0.3 - 0.6) |
| ln (monocytes, ×10<sup>3</sup>/μL) | 2872 | -0.826 ± 0.377 |
| Neutrophils, ×10<sup>3</sup>/μL | 2873 | 3.4 (2.7 - 4.3) |
| ln (neutrophils, ×10<sup>3</sup>/μL) | 2873 | 1.21 ± 0.39 |
| Total white blood cells, ×10<sup>3</sup>/μL | 2873 | 5.8 (4.8 – 7.0) |
| ln (total white blood cells, ×10<sup>3</sup>/μL) | 2873 | 1.76 ± 0.29 |
| Red blood cells, ×10<sup>6</sup>/μL | 2873 | 4.48 ± 0.49 |
| Platelets, ×10<sup>3</sup>/μL | 2868 | 228.4 ± 62.5 |

Data are expressed as mean ± SD, n (%), or median (interquartile range). MET, metabolic equivalent.

### Association of total cholesterol levels with cell counts

In the fully adjusted model, a higher total cholesterol level was associated with a lower ln-transformed total white blood cell count after adjusting for multiple testing correction (B=-0.028, *p*<0.001 with a 2.8% lower cell count per one SD increase in total cholesterol level (Supplementary Table 2 and Figure 1). When assessing different white blood cell types, a higher total cholesterol level was also significantly associated with lower ln-transformed monocyte count (B=-0.038, *p*=<0.001) and neutrophil count (B=-0.034, *p*=<0.001), with 3.7% and 3.4% lower cell count, respectively per one SD increase in total cholesterol level after multiple testing correction (Supplementary Table 2 and Figure 1). In the sensitivity analysis, whereby participants taking lipid-lowering medications were excluded, a higher total cholesterol level was also significantly associated with lower ln-transformed total white blood cell, monocyte and neutrophil counts (*p*=0.004, 0.006, and 0.009 respectively, Figure 2). In all the above analyses, no significant interaction with sex and race/ethnicity was observed.

**Figure 1.**
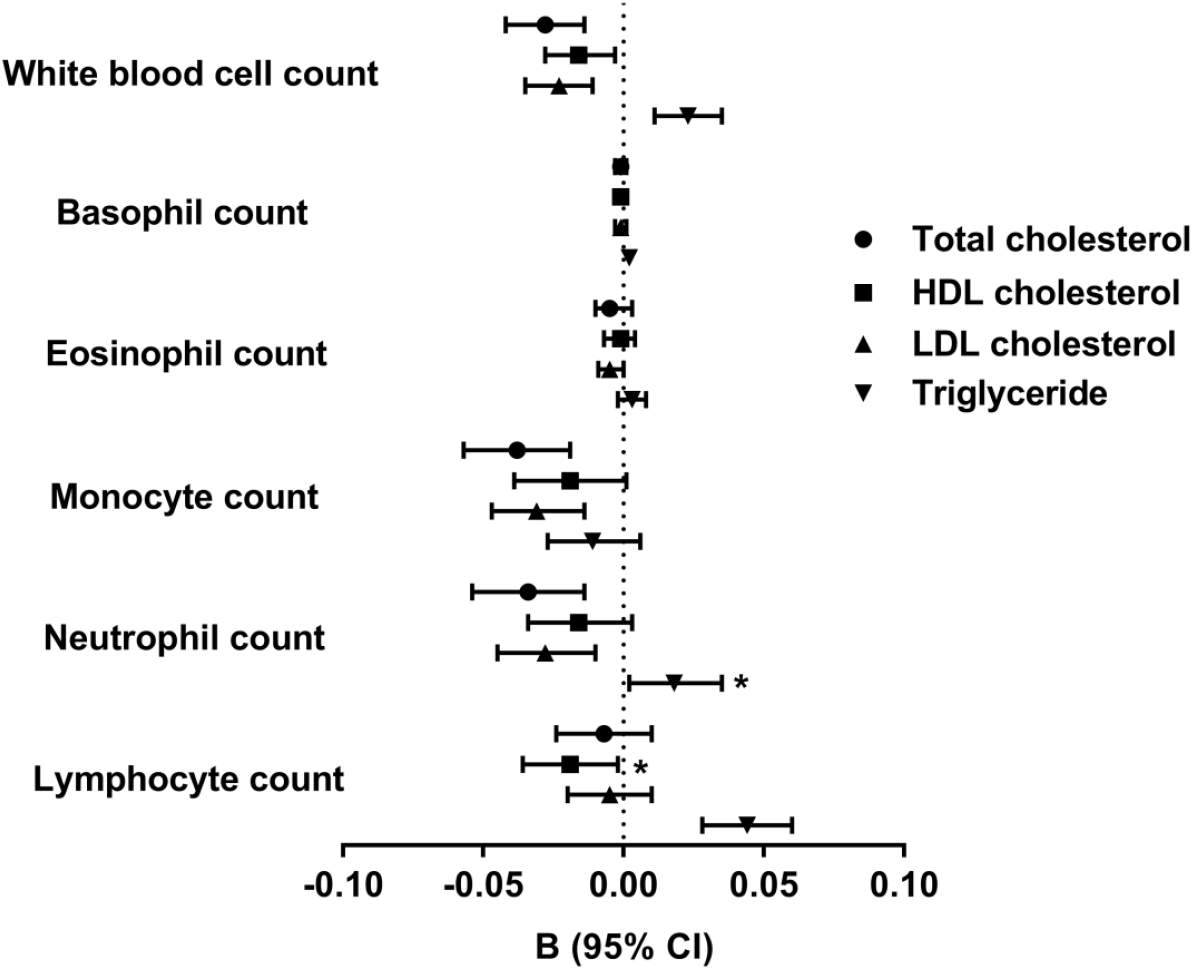
Association of lipid levels with different types of white blood cell counts. Regression coefficient B (95% confidence interval) were expressed as ln-transformed cell counts (×10^3^/μL) per one SD unit increase in lipid levels. Data on different white blood cell counts and triglycerides were ln-transformed in the analysis. Data were all adjusted for age, sex, ethnicity, BMI, family income, smoking, pack-years of smoking, current alcohol use, physical activity, hypertension, diabetes, use of any lipid-lowering medication, eGFR, *APOE* genotype, history of CVD, history of coronary revascularization, red blood cell and platelet counts. Data were also adjusted for ln-transformed triglycerides and HDL cholesterol for the analysis of total cholesterol and LDL cholesterol; LDL cholesterol and ln-transformed triglycerides for analysis of HDL cholesterol; and HDL cholesterol and LDL cholesterol for analysis of ln-transformed triglycerides. Asterisks (*) indicate *P* values that did not remain significant after multiple testing correction.

**Figure 2.**
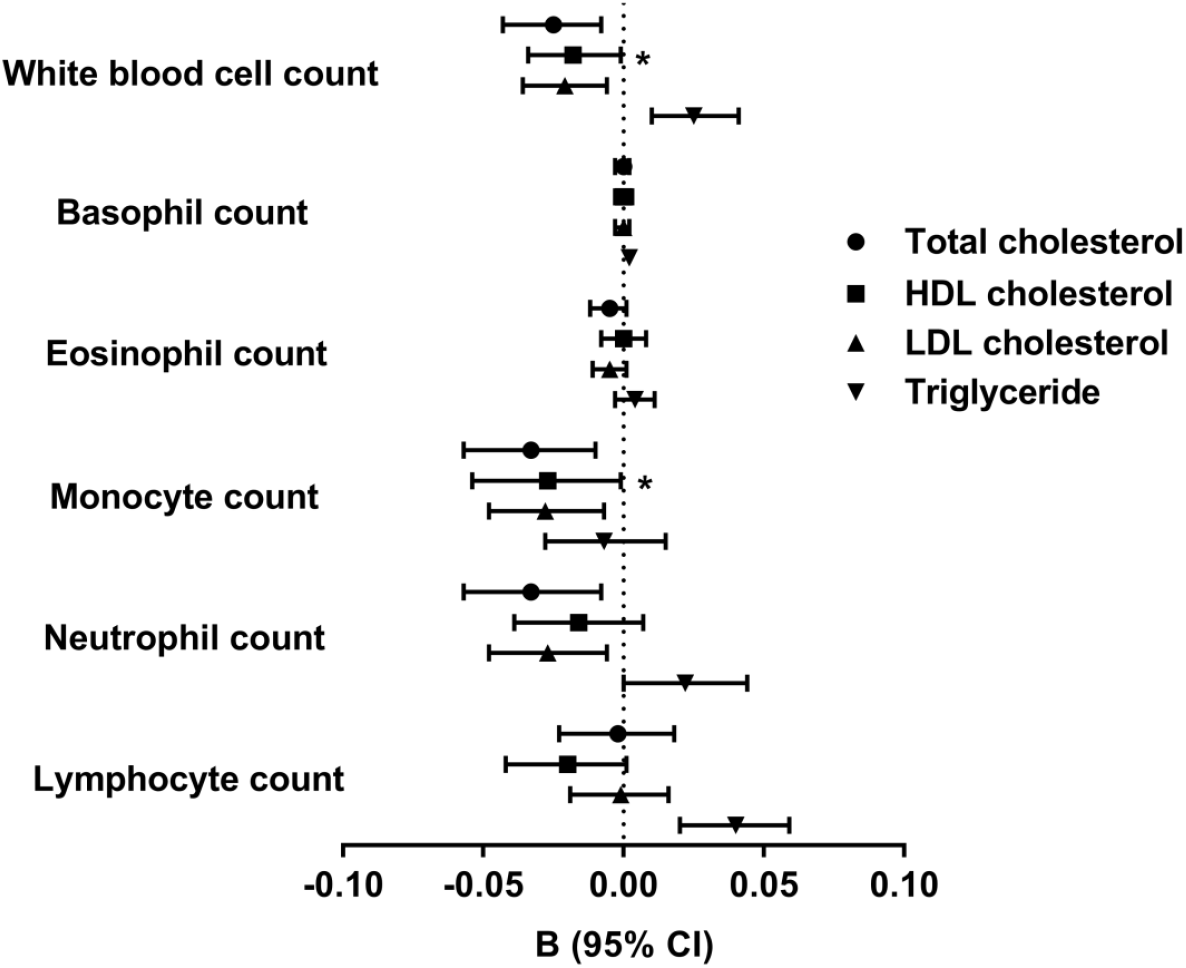
Association of lipid levels with different types of white blood cell counts after excluding participants taking lipid-lowering drugs (n=1715). Regression coefficient B (95% confidence interval) were expressed as ln-transformed cell counts (×10^3^/μL) per one SD unit increase in lipid levels. Data on different white blood cell counts and triglycerides were ln-transformed in the analysis. All data were adjusted for age, sex, ethnicity, BMI, family income, smoking, pack-years of smoking, current alcohol use, physical activity, hypertension, diabetes, eGFR, *APOE* genotype, history of CVD, history of coronary revascularization, red blood cell counts and platelet counts. Data were also adjusted for ln-transformed triglycerides and HDL cholesterol for analysis of total cholesterol and LDL cholesterol; LDL cholesterol and ln-transformed triglycerides for analysis of HDL cholesterol; and HDL cholesterol and LDL cholesterol for analysis of ln-transformed triglycerides. Asterisks (*) indicate *P* values that did not remain significant after multiple testing correction.

In a separate analysis, we assessed the non-linear association of total cholesterol levels with total white blood cell and neutrophil counts. The regression spline analysis suggested that both the associations may be somewhat steeper at lower total cholesterol levels, and flatter at higher total cholesterol levels, with a threshold at approximately 155 and 204 mg/dL, respectively (Table 2). However, such non-linear associations were not found after excluding participants taking lipid-lowering medications. No non-linear association was found for the different types of white blood cell counts.

**Table 2.** Non-linear association of different lipid levels with ln-transformed white blood cell counts

| **Lipid measures** | **n** | **B (95% CI)** | **p-value** |
| --- | --- | --- | --- |
| **White blood cell count** |  |  |  |
| Total cholesterol |  |  |  |
| <155 mg/dL | 708 | -0.053 (-0.120, 0.015) | 0.124 |
| ≥155 mg/dL | 2165 | -0.019 (-0.036, -0.002) | 0.031 |
| LDL cholesterol |  |  |  |
| <125 mg/dL | 2137 | -0.038 (-0.059, -0.017) | <0.001 |
| ≥125 mg/dL | 736 | 0.029 (-0.004, -0.061) | 0.086 |
| **Neutrophil count** |  |  |  |
| Total cholesterol |  |  |  |
| <204 mg/dL | 2136 | -0.057 (-0.089, -0.024) | <0.001 |
| ≥204 mg/dL | 737 | 0.041 (-0.008, 0.091) | 0.103 |

Regression coefficient B (95% confidence interval) were expressed as ln-transformed cell counts
( $\times 10^3/\mu\text{L}$ ). All data were adjusted for age, sex, ethnicity, BMI, family income, smoking, pack-years of smoking, current alcohol use, physical activity, hypertension, diabetes, use of any lipid-lowering medication, eGFR, *APOE* genotype, history of CVD, history of coronary revascularization, ln-transformed triglycerides, HDL cholesterol, red blood cell counts and platelet counts.

In all above analyses, similar non-significant results after multiple testing correction were obtained when assessing basophil and eosinophil counts without ln-transformation (Supplementary Figures 2 and 3).

### Association of HDL cholesterol levels with cell counts

A higher HDL cholesterol level was significantly associated with a lower ln-transformed total white blood cell count in the full adjustment model (B=-0.016, *p*=0.018), with a 1.5% lower count per one SD increase in HDL cholesterol level after correcting for multiple comparisons (Supplementary Table 2 and Figure 1). However, such association did not remain significant in the sensitivity analysis after excluding participants taking lipid-lowering medications (Figure 2). When analysing different white blood cell types, no significant association was found after full adjustment and multiple testing correction (Supplementary Table 2, and Figures 1 & 2). In all the above analyses, no significant interaction with sex and race/ethnicity was found.

Similar non-significant results were obtained when assessing basophil and eosinophil counts without ln-transformation (Supplementary Figures 2 and 3). No non-linear association was found with total white blood cell counts and different types of white blood cell counts.

### Association of LDL cholesterol levels with cell counts

A higher LDL cholesterol level was significantly associated with a lower ln-transformed total white blood cell count in the full adjustment model (B=-0.023, *p*<0.001), with a 2.3% lower cell count per one SD increase in LDL cholesterol level after multiple testing correction (Supplementary Table 2 and Figure 1). When assessing different white blood cell types, a higher LDL cholesterol level was also significantly associated with a lower ln-transformed monocyte count (B=-0.031, *p*<0.001) and neutrophil count (B=-0.028, *p*=0.002) with 3.0% and 2.7% lower cell counts, respectively per one SD increase in LDL cholesterol level in the full adjustment model after multiple testing correction. In a sensitivity analysis after excluding participants taking lipid-lowering drugs, a higher LDL cholesterol level was still significantly associated with a lower ln-transformed total white blood cell, monocyte and neutrophil counts (*p*=0.006, 0.007, and 0.013 respectively, Figure 2). In all the above analyses, no significant interaction with sex and race/ethnicity was found.

In a separate analysis, we assessed the non-linear association of LDL cholesterol levels with total white blood cell counts. The regression spline analysis suggested that the association may be somewhat steeper at lower LDL cholesterol levels, and flatter at higher LDL cholesterol levels, with a threshold at approximately 125 mg/dL (Table 2). However, such non-linear association was not found after excluding participants taking lipid-lowering medications. No non-linear association was found for the different types of white blood cell counts.

In all above analyses, similar non-significant results were obtained after multiple testing correction when assessing basophil and eosinophil counts without ln-transformation. (Supplementary Figures 2 and 3).

### Association of triglyceride levels with cell counts

A higher ln-transformed triglyceride level was significantly associated with a higher ln-transformed total white blood cell count in the full adjustment model after multiple testing correction (β=0.023, *p*<0.001) with a 2.3% higher cell count per one SD increase in ln-transformed triglyceride level (Supplementary Table 2 and Figure 1). A significant interaction with race/ethnicity was found (*p*=0.005), but not with sex. The association was significant only in African Americans (*p*<0.001), but not in non-Hispanic white Americans and Hispanic Americans (Supplementary Table 3). When assessing different types of white blood cell counts, a higher ln-transformed triglyceride level was associated with a higher ln-transformed lymphocyte count (B=0.044, *p*<0.001) in the full adjustment model after multiple testing correction (Supplementary Table 2 and Figure 1), with a 4.5% higher lymphocyte count per one SD increase in ln-transformed triglyceride level. No significant interaction with sex and race/ethnicity was found. No significant association was found between triglyceride levels and other types of white blood cell counts after multiple testing correction.

In the sensitivity analysis after excluding participants taking lipid-lowering drugs, a higher ln-transformed triglyceride level was still significantly associated with a higher ln-transformed total white blood cell count and a higher lymphocyte count after multiple testing correction (*p*=0.001 and <0.001 respectively, Figure 2). No significant interaction with sex or race/ethnicity was found.

In all above analyses, similar non-significant results were obtained when assessing basophil and eosinophil counts without ln-transformation (Supplementary Figures 2 and 3). No non-linear association was found with total white blood cell counts and different types of white blood cell counts.

### Inflammation and infection related conditions as potential confounders

Since white blood cell counts can be elevated in inflammatory and infection related conditions, a sensitivity analysis was performed after excluding 1069 participants with these conditions. Similar to the main analysis, higher total and LDL cholesterol levels were associated with lower ln-transformed total white blood cell, monocyte, and neutrophil counts, whereas higher triglyceride levels were associated with higher ln-transformed total white blood cell and lymphocyte counts (Supplementary Figure 4). Interestingly, higher HDL cholesterol levels were found to be associated with a lower ln-transformed total white blood cell counts in this sensitivity analysis (p = 0.007, Supplementary Figure 4), a relationship that was previously observed in the main cohort (Figure 1), but not in the sensitivity analysis after excluding participants taking lipid-lowering medication (Figure 2). Similar non-significant results were obtained when assessing basophil and eosinophil counts without ln-transformation (Supplementary Figure 5).

## Discussion

Using data from MESA, this study examined the relationships between various plasma lipids and a range of white blood cell counts in a multi-ethnic cohort. After adjusting for relevant co-variates, higher total cholesterol and LDL cholesterol levels were significantly associated with lower white blood cell, monocyte and neutrophil counts in all participants. Higher triglyceride levels were associated with higher total white blood cell and lymphocyte counts. Increased HDL cholesterol was associated with lower total white blood cell counts, but the association did not remain significant after excluding participants taking lipid-lowering medication. Despite these significant associations, the magnitudes of these associations were modest, with a one standard deviation change in lipid levels resulting in less than 5% change in corresponding white blood cell counts. The results of this study suggest that relationships between plasma lipid levels and white blood cell counts are not common amongst mice and humans.

Elevated cholesterol levels have been demonstrated to promote monocytosis in mice due to cholesterol accumulation and proliferation of hematopoietic stem progenitor cells (HSPCs)^2, 4, 23^. Recently, van der Valk, et al.^24^ reported that cholesterol accumulation can occur in human HSPCs and results in a myeloid differentiation bias *in vitro.* Whilst that study also reported increased glucose uptake in the bone marrow of subjects with familial hypercholesterolemia, suggesting increased bone marrow progenitor proliferation, monocyte, lymphocyte and neutrophil counts were not elevated *in* vivo. In light of the findings presented here, further investigation into the impact of cholesterol accumulation on the mechanisms underlying myeloid cell differentiation is needed.

Neutrophils play an important role in atherosclerosis. They accumulate preferentially in the shoulder regions of atherosclerotic plaques in mice, where there is also a high density of monocytes^3, 25, 26^. This is consistent with atherosclerotic lesion size in mice being positively correlated with neutrophil counts, suggesting a potential role of neutrophils in atherosclerotic lesion development^3^. Indeed, several studies have shown that hypercholesterolaemia induces neutrophilia in mice^2, 27, 28^. Similarly, a cross-sectional analysis by Huang, et al.^29^ looking at the effect of hyperlipidaemia, smoking and BMI on peripheral differential leukocyte counts in humans also showed a positive correlation between LDL cholesterol and neutrophil counts. However, we found that elevated total cholesterol and LDL was associated with a significant, if modest, decrease in neutrophil counts. Indeed, children with familial hypercholesterolaemia have normal neutrophil counts despite their high cholesterol levels^4^.

The effect of triglycerides on CVD risk is well established^30, 31^. Increased bone marrow glucose uptake, together with monocytosis and leukocytosis has been described in patients with familial dysbetalipoproteinaemia^6^, and increased triglyceride rich remnant cholesterol has been associated with increased total white blood cell, monocyte, neutrophil and lymphocyte counts in a large population^6^. Our analysis found an association of higher triglyceride levels with higher total white blood cell and lymphocyte counts similar to those previously described by Oda and colleagues^7,8^, although the same relationship between LDL and lymphocyte number was not observed. Elevated circulating lymphocyte numbers may be due to increased lipid loaded, antigen presenting arterial wall macrophages^7^. However, as lymphocytes comprise a heterogenous group of cells, further studies are needed to identify which type of lymphocytes are affected by elevated triglyceride levels.

Our analysis found that monocyte counts were not associated with higher plasma triglyceride levels. Saja, et al.^32^ demonstrated that in a murine model of hypertriglyceridaemia increased binding of Gr1^low^ monocytes to the endothelium and extravasation into the subendothelial space. This resulted in depletion of monocytes from the circulation and accumulation of macrophages in the heart, liver, and kidney. In this context, it may be important to determine the if there is any relationship between plasma lipid levels and monocyte flux in humans, as the number of monocytes in the blood represents only a snapshot of the balance between the emergence of cells from the bone marrow and migration of monocytes into peripheral tissues.

Inflammation and infection result in the elevation of different white blood cell counts, and may therefore have confounding effect on the relationship between lipids and white blood cell counts. However, in the present analysis, excluding participants with inflammation and infection related diseases did not have any significant impact on the relationships observed between lipid levels and white blood cell counts. Future analyses could utilise datasets that include reliable direct measures of inflammation (such as CRP). This data was not available for a sufficient number of subjects in visit 5 of the MESA study but may be important in the context of HDL function that is known to be impaired under inflammatory conditions^33^, and persists for a significant period of time after the inflammatory event^34^.

Whilst the present study takes the advantage of a large well-characterised sample of with a multi-ethnic study design, there are some limitations. We were not able to definitively determine the familial hypercholesterolemia (FH) status of subjects included in our study. Future studies using larger populations with data on genetic mutations and/or relevant patient history may allow for more accurate status determination and assessment of the impact of FH on white blood cell counts. There was an under-representation of Chinese Americans in the present study, compared to the original parent cohort. Therefore, we could not assess ethnic differences for Chinese Americans. White blood cell profiles were only measured at exam 5 and not at baseline exam 1. As such, we could not assess the temporal relationship between changes in lipid measures and change in white blood cell counts. The current analysis was not able to consider the potential confounding effect of medications that may affect white blood cell counts and inflammation status. As this was an observational study, further investigations are needed to determine the underlying mechanism of the associations and their clinical significance. Nevertheless, this is the largest study to date with multi-ethnic study design that has assessed the relationship between lipid measures and different white blood cell counts.

In conclusion, this study suggests modest associations between lipid levels and circulating leukocyte counts in humans. Further studies in other clinical settings are needed to confirm our findings. Whilst significant relationships were observed, the relative contribution of lipids and inflammatory factors in driving atherosclerosis in humans remains an open question.

## Acknowledgements

Kwok Leung Ong was supported by an Australian National Health and Medical Research Council Career Development Fellowship (1122854). Blake J. Cochran was supported by an International Atherosclerosis Society Fellowship. The MESA study was supported by contracts HHSN268201500003I, N01-HC-95159, N01-HC-95160, N01-HC-95161, N01-HC-95162, N01-HC-95163, N01-HC-95164, N01-HC-95165, N01-HC-95166, N01-HC-95167, N01-HC-95168 and N01-HC-95169 from the National Heart, Lung, and Blood Institute, by grants UL1-TR-000040, UL1-TR-001079, and UL1-TR-001420 from National Center for Advancing Translational Sciences. The authors thank the other investigators, the staff, and the participants of the MESA study for their valuable contributions. A full list of participating MESA investigators and institutions can be found at http://www.mesa-nhlbi.org.

## Contribution statement

KLO and BJC conceived the study. YCL, RLM, MAA and KLO designed the statistical analysis. YCL, KLO and BJC performed the data analysis and interpretation, with assistance and guidance from KLW, RLM, MAA and KAR. YCL and KLO completed all analyses. YCL, KLO and BJC wrote the manuscript with input and critical insights from KJW, RLM, MAA and KAR. All authors have approved the final manuscript.

---

**Supplementary Table 1.** Clinical characteristics of participants included and not included in this

| Characteristics | Not included | Included | *p*-value |
| --- | --- | --- | --- |
| n | 1843 | 2873 |  |
| Age, years | 70.5 ± 9.7 | 69.5 ± 9.3 | <0.001 |
| Women | 995 (54.0%) | 1519 (52.9%) | 0.453 |
| Race/ethnicity |  |  | <0.001 |
| Non-Hispanic White | 695 (37.7%) | 1231 (42.8%) |  |
| African American | 447 (24.3%) | 803 (27.9%) |  |
| Hispanic American | 187 (10.1%) | 812 (28.3%) |  |
| Chinese American | 514 (27.9%) | 27 (0.9%) |  |
| Recruitment site |  |  | <0.001 |
| Forsyth County, North Carolina | 215 (11.7%) | 598 (20.8%) |  |
| New York, New York | 148 (8.0%) | 662 (23.0%) |  |
| Baltimore, Maryland | 154 (8.4%) | 504 (17.5%) |  |
| St. Paul, Minnesota | 77 (4.2%) | 694 (24.2%) |  |
| Chicago, Illinois | 833 (45.2%) | 43 (1.5%) |  |
| Los Angeles County, California | 416 (22.6%) | 372 (12.9%) |  |

Data are expressed as mean ± SD or n (%). Baseline characteristics were determined using a chi-square test for categorical variables, and an ANOVA test for continuous variables.

**Supplementary Table 2.**
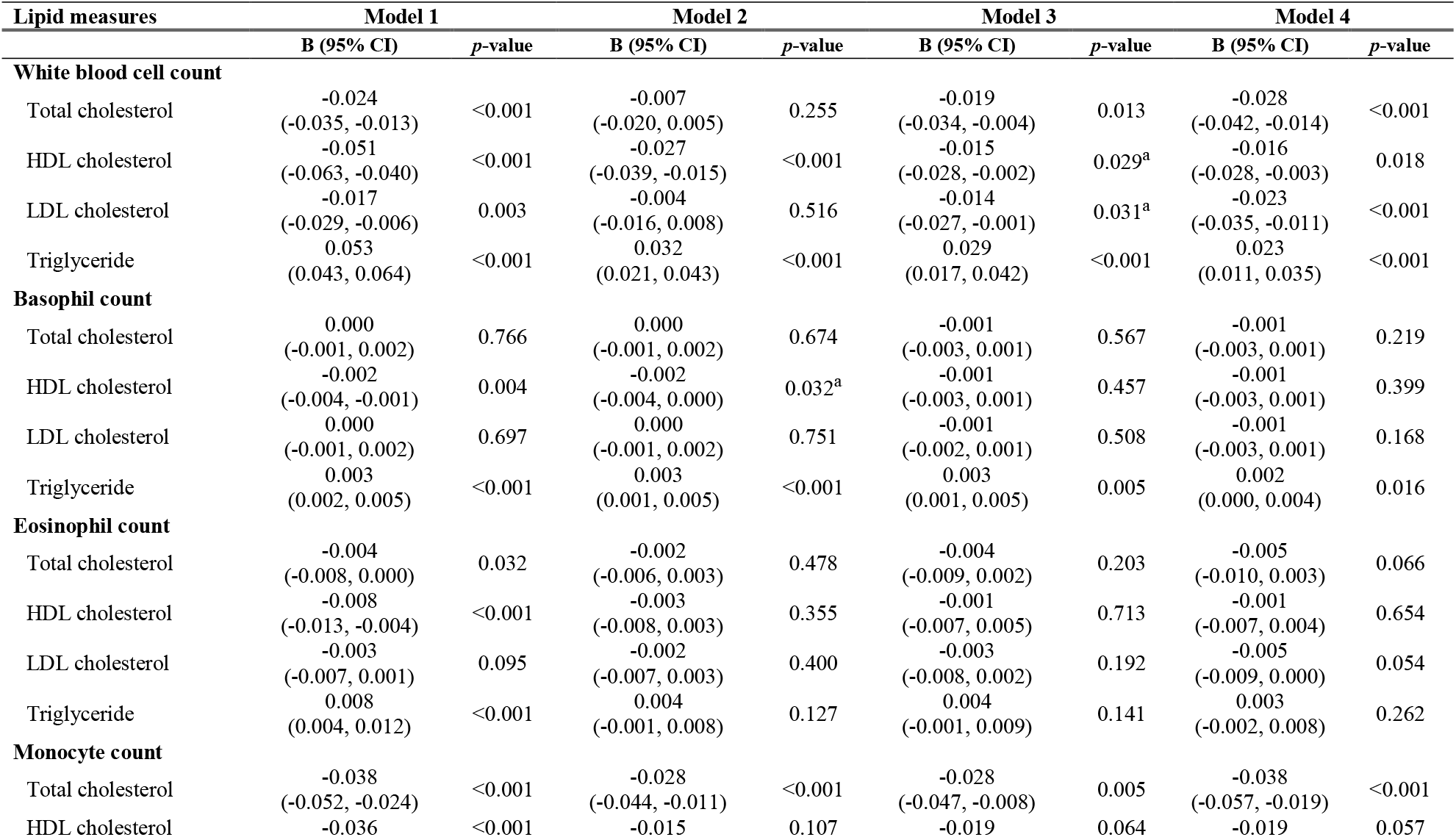
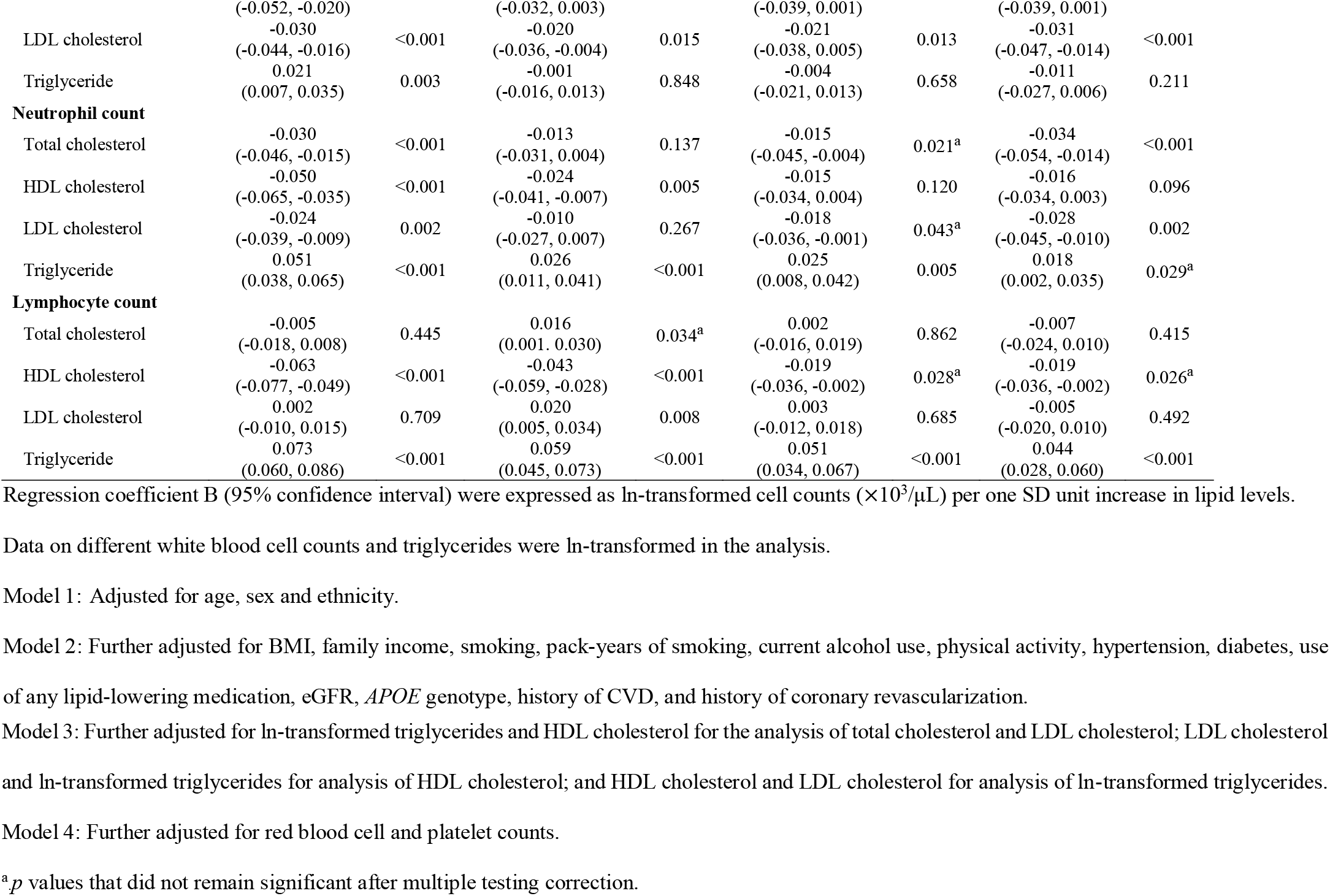
Association of lipid levels with different types of white blood cell counts

**Supplementary Table 3.**
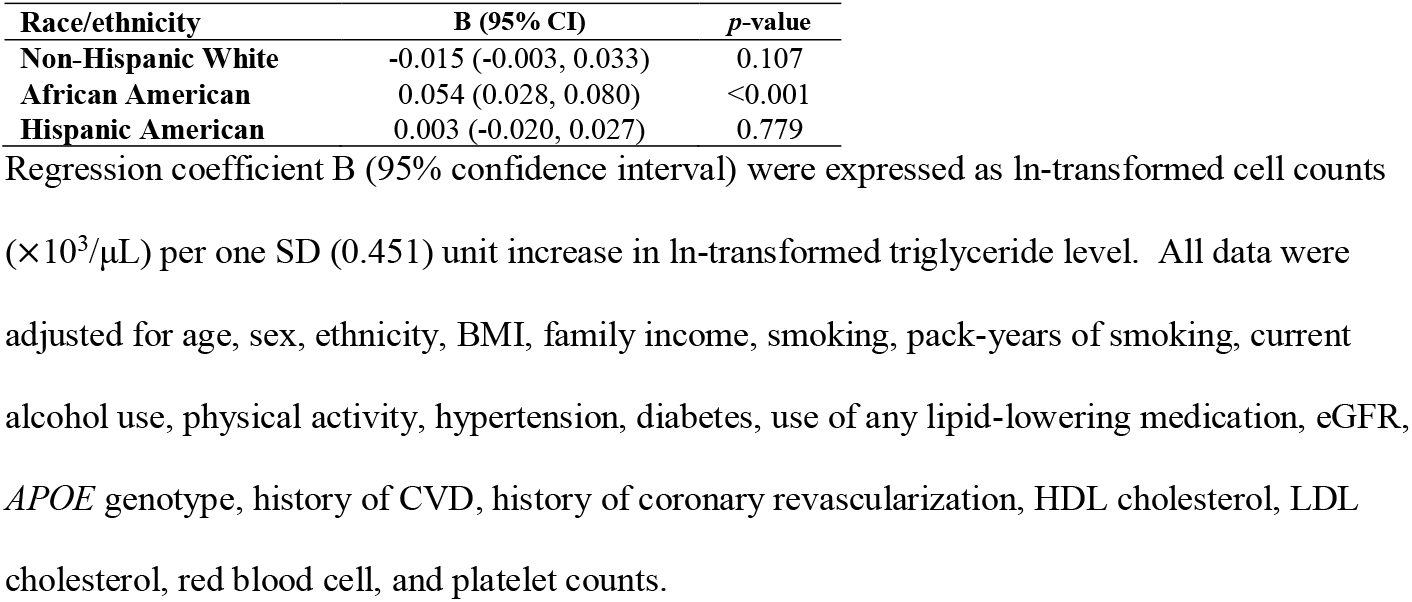
Association of triglycerides levels with total white blood cell counts according to race/ethnicity using multivariable linear regression analysis (n=2873)

| **Race/ethnicity** | **B (95% CI)** | **p-value** |
| --- | --- | --- |
| **Non-Hispanic White** | -0.015 (-0.003, 0.033) | 0.107 |
| **African American** | 0.054 (0.028, 0.080) | <0.001 |
| **Hispanic American** | 0.003 (-0.020, 0.027) | 0.779 |

Regression coefficient B (95% confidence interval) were expressed as ln-transformed cell counts ( $\times 10^3/\mu\text{L}$ ) per one SD (0.451) unit increase in ln-transformed triglyceride level. All data were adjusted for age, sex, ethnicity, BMI, family income, smoking, pack-years of smoking, current alcohol use, physical activity, hypertension, diabetes, use of any lipid-lowering medication, eGFR, *APOE* genotype, history of CVD, history of coronary revascularization, HDL cholesterol, LDL cholesterol, red blood cell, and platelet counts.

**Supplementary Figure 1.**
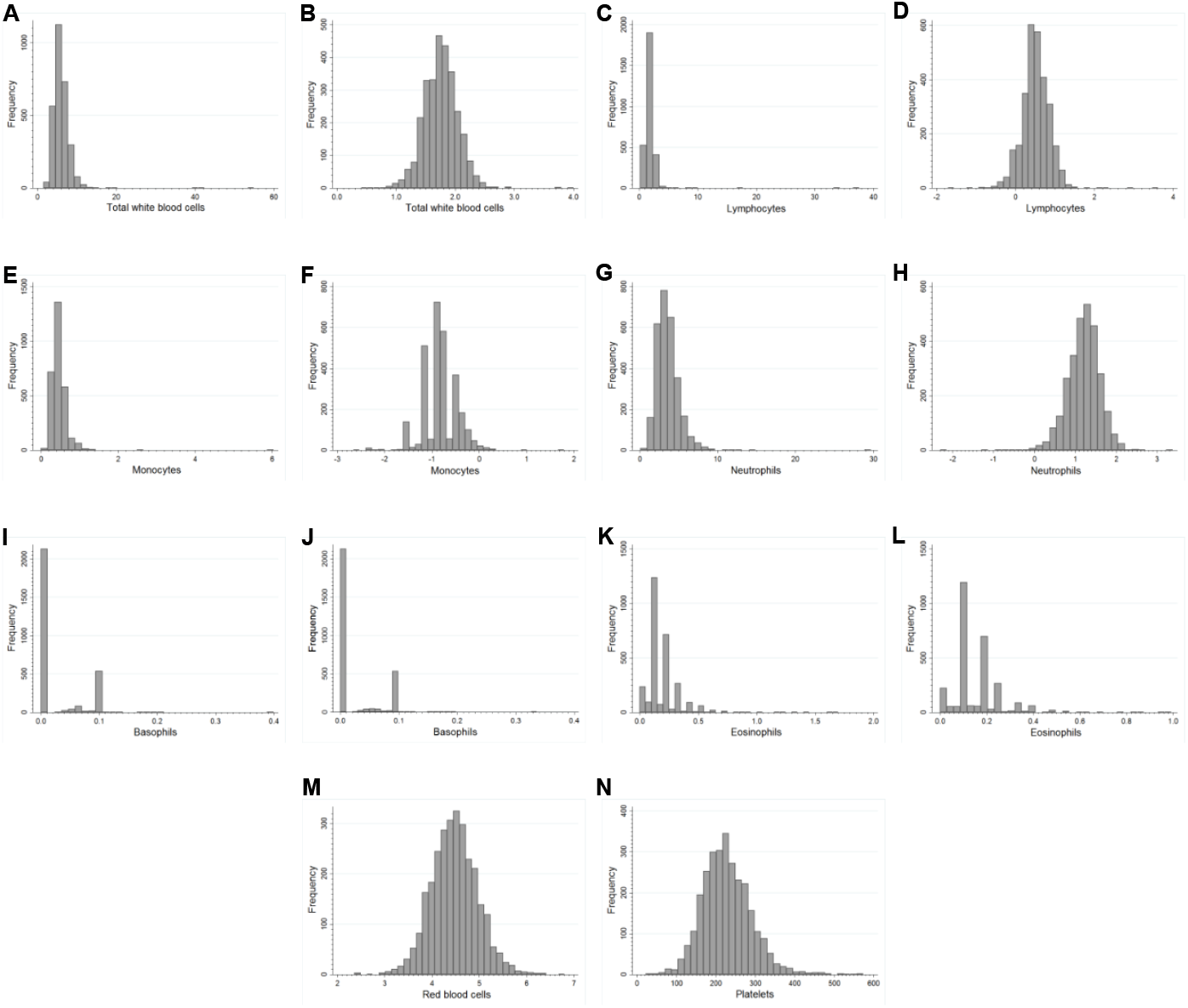
Distribution of total white blood cell (A, B), lymphocyte (C, D), monocyte (E, F), neutrophil (G, H), basophil (I, J) and eosinophil (K, L) counts (×10^3^/μL), before (A, C, E, G, I and K) and after (B, D, F, H, J and L) ln-transformation. Distribution of untransformed red blood cell (M) (×10^6^/μL) and platelet (N) (×10^3^/μL) counts.

**Supplementary Figure 2.**
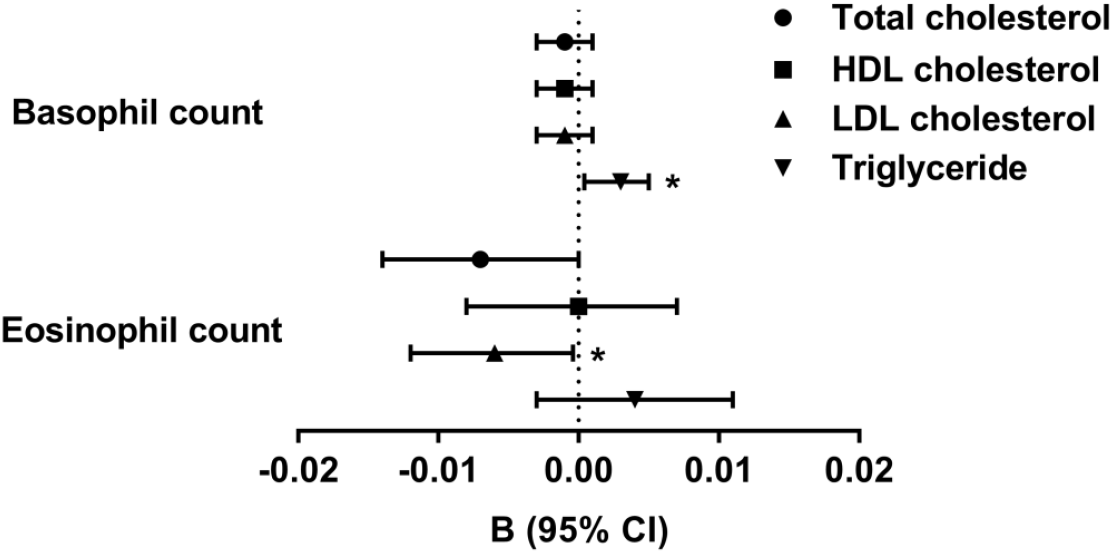
Association of lipid levels with un-transformed basophil and eosinophil counts using multivariable linear regression analysis (n=2873). Regression coefficient B (95% confidence interval) were expressed as cell counts (×10^3^/μL) per one SD unit increase in lipid levels. Data on triglycerides were ln-transformed in the analysis. All data were adjusted for age, sex, ethnicity, BMI, family income, smoking, pack-years of smoking, current alcohol use, physical activity, hypertension, diabetes, use of any lipid-lowering medication, eGFR, *APOE* genotype, history of CVD, history of coronary revascularization, red blood cell and platelet counts. Data were also adjusted for ln-transformed triglycerides and HDL cholesterol for the analysis of total cholesterol and LDL cholesterol; LDL cholesterol and ln-transformed triglycerides for analysis of HDL cholesterol; and HDL cholesterol and LDL cholesterol for analysis of ln-transformed triglycerides.

**Supplementary Figure 3.**
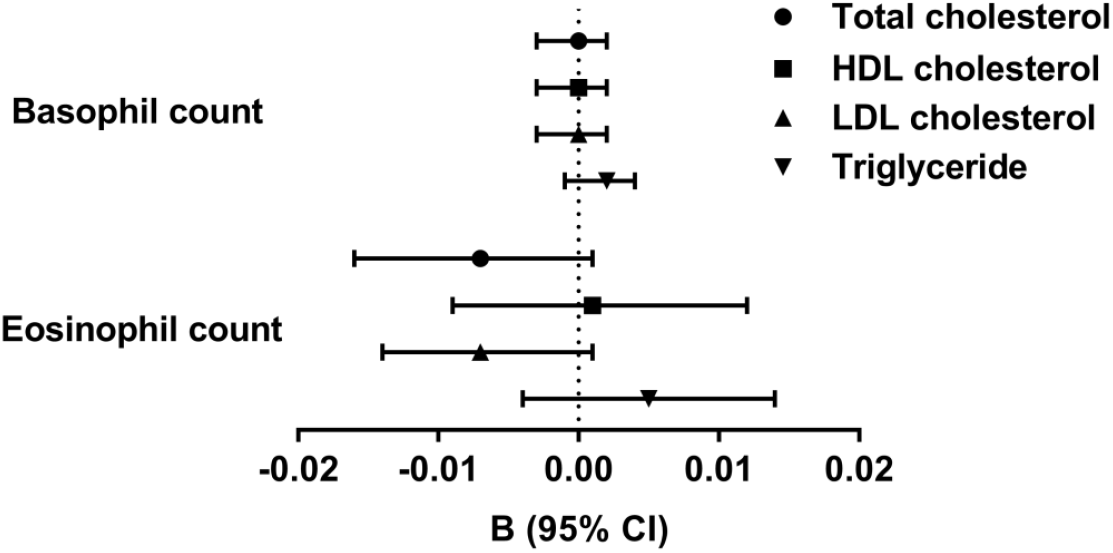
Association of lipid levels with un-transformed basophil and eosinophil counts using multivariable linear regression analysis after excluding participants taking lipid-lowering drugs (n=1715). Regression coefficient B (95% confidence interval) were expressed as cell counts (×10^3^/μL) per one SD unit increase in lipid levels. Data on triglycerides were ln-transformed in the analysis. All data were adjusted for age, sex, ethnicity, BMI, family income, smoking, pack-years of smoking, current alcohol use, physical activity, hypertension, diabetes, eGFR, *APOE* genotype, history of CVD, history of coronary revascularization, red blood cell counts and platelet counts. Data were also adjusted for ln-transformed triglycerides and HDL cholesterol for analysis of total cholesterol and LDL cholesterol; LDL cholesterol and ln-transformed triglycerides for analysis of HDL cholesterol; and HDL cholesterol and LDL cholesterol for analysis of ln-transformed triglycerides. Asterisks (*) indicate *P* values that did not remain significant after multiple testing correction.

**Supplementary Figure 4.**
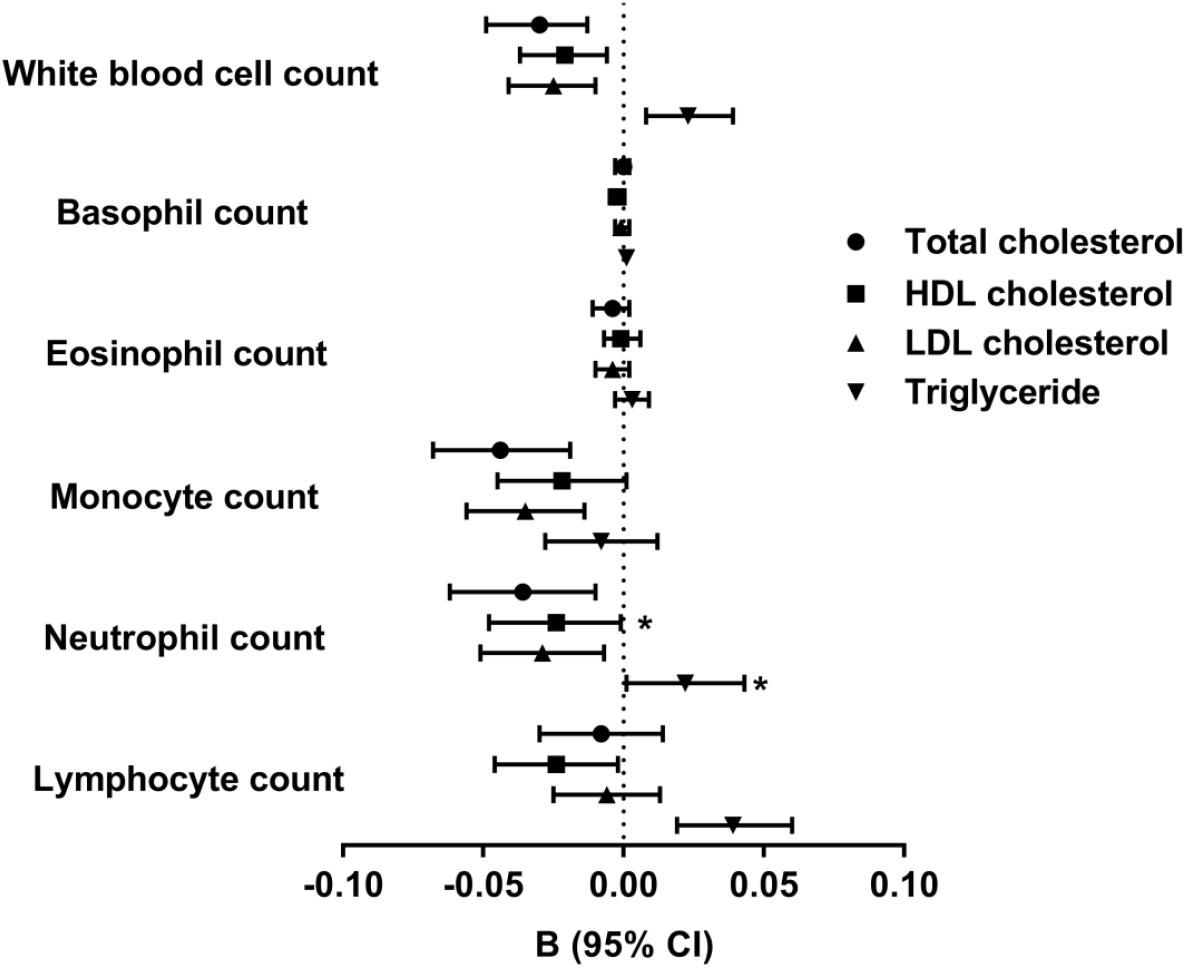
Association of lipid levels with different types of white blood cell counts among participants without self-reported confounding disease (n=1804). Participants were included in this analysis if they reported not having any fever, cold/flu, urinary infection, seasonal allergy, bronchitis, sinus infection, pneumonia, tooth infection or liver problem over the past two weeks in the medical history questionnaires. Regression coefficient B (95% confidence interval) were expressed as ln-transformed cell counts (×10^3^/μL) per one SD unit increase in lipid levels. Data on different white blood cell counts and triglycerides were ln-transformed in the analysis. All data were adjusted for age, sex, ethnicity, BMI, family income, smoking, pack-years of smoking, current alcohol use, physical activity, hypertension, diabetes, eGFR, *APOE* genotype, history of CVD, history of coronary revascularization, red blood cell counts and platelet counts. Data were also adjusted for ln-transformed triglycerides and HDL cholesterol for analysis of total cholesterol and LDL cholesterol; LDL cholesterol and ln-transformed triglycerides for analysis of HDL cholesterol; and HDL cholesterol and LDL cholesterol for analysis of ln-transformed triglycerides. Asterisks (*) indicate *P* values that did not remain significant after multiple testing correction.

**Supplementary Figure 5.**
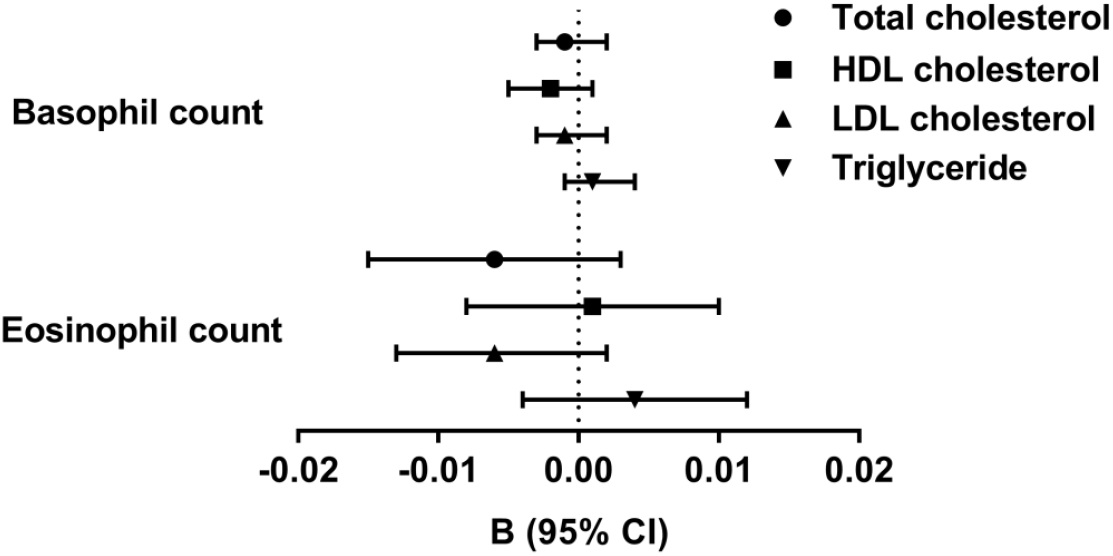
Association of lipid levels with un-transformed basophil and eosinophil counts using multivariable linear regression analysis among participants without self-reported confounding disease (n=1804). Participants were included in this analysis if they reported not having any fever, cold/flu, urinary infection, seasonal allergy, bronchitis, sinus infection, pneumonia, tooth infection or liver problem over the past two weeks in the medical history questionnaires. Regression coefficient B (95% confidence interval) were expressed as cell counts (×10^3^/μL) per one SD unit increase in lipid levels. Data on triglycerides were ln-transformed in the analysis. All data were adjusted for age, sex, ethnicity, body mass index, family income, smoking, pack-years of smoking, current alcohol use, physical activity, hypertension, diabetes, eGFR, *APOE* genotype, history of CVD, history of coronary revascularization, red blood cell counts and platelet counts. Data were also adjusted for ln-transformed triglycerides and HDL cholesterol for analysis of total cholesterol and LDL cholesterol; LDL cholesterol and ln-transformed triglycerides for analysis of HDL cholesterol; and HDL cholesterol and LDL cholesterol for analysis of ln-transformed triglycerides.

## References

1. Koltsova EK, Hedrick CC, Ley K. Myeloid cells in atherosclerosis: A delicate balance of antiinflammatory and proinflammatory mechanisms. Curr Opin Lipidol. 2013;24:371–380

2. Murphy AJ, Akhtari M, Tolani S, Pagler T, Bijl N, Kuo C-L, Wang M, Sanson M, Abramowicz S, Welch C, Bochem AE, Kuivenhoven JA, Yvan-Charvet L, Tall AR. Apoe regulates hematopoietic stem cell proliferation, monocytosis, and monocyte accumulation in atherosclerotic lesions in mice. The Journal of Clinical Investigation. 2011;121:4138–4149

3. Drechsler M, Megens RTA, van Zandvoort M, Weber C, Soehnlein O. Hyperlipidemia-triggered neutrophilia promotes early atherosclerosis. Circulation. 2010;122:1837

4. Tolani S, Pagler TA, Murphy AJ, Bochem AE, Abramowicz S, Welch C, Nagareddy PR, Holleran S, Hovingh GK, Kuivenhoven JA, Tall AR. Hypercholesterolemia and reduced hdl-c promote hematopoietic stem cell proliferation and monocytosis: Studies in mice and fh children. Atherosclerosis. 2013;229:79–85

5. Andersen CJ, Vance TM. Gender dictates the relationship between serum lipids and leukocyte counts in the national health and nutrition examination survey 1999(-)2004. J Clin Med. 2019;8

6. Bernelot Moens SJ, Verweij SL, Schnitzler JG, Stiekema LCA, Bos M, Langsted A, Kuijk C, Bekkering S, Voermans C, Verberne HJ, Nordestgaard BG, Stroes ESG, Kroon J. Remnant cholesterol elicits arterial wall inflammation and a multilevel cellular immune response in humans. Arteriosclerosis, thrombosis, and vascular biology. 2017;37:969–975

7. Oda E. Longitudinal associations between lymphocyte count and ldl cholesterol in a health screening population. J Clin Transl Endocrinol. 2014;1:49–53

8. Oda E, Kawai R, Aizawa Y. Lymphocyte count was significantly associated with hyper-ldl cholesterolemia independently of high-sensitivity c-reactive protein in apparently healthy japanese. Heart and vessels. 2012;27:377–383

9. Fessler MB, Rose K, Zhang Y, Jaramillo R, Zeldin DC. Relationship between serum cholesterol and indices of erythrocytes and platelets in the us population. Journal of Lipid Research. 2013;54:3177–3188

10. Rothe G, Gabriel H, Kovacs E, Klucken J, Stohr J, Kindermann W, Schmitz G. Peripheral blood mononuclear phagocyte subpopulations as cellular markers in hypercholesterolemia. Arterioscler Thromb Vasc Biol. 1996;16:1437–1447

11. Armstrong AJ, Gebre AK, Parks JS, Hedrick CC. Atp-binding cassette transporter g1 negatively regulates thymocyte and peripheral lymphocyte proliferation. J Immunol. 2010;184:173–183

12. Bensinger SJ, Bradley MN, Joseph SB, Zelcer N, Janssen EM, Hausner MA, Shih R, Parks JS, Edwards PA, Jamieson BD, Tontonoz P. Lxr signaling couples sterol metabolism to proliferation in the acquired immune response. Cell. 2008;134:97–111

13. Surls J, Nazarov-Stoica C, Kehl M, Olsen C, Casares S, Brumeanu TD. Increased membrane cholesterol in lymphocytes diverts t-cells toward an inflammatory response. PLoS One. 2012;7:e38733

14. Bild DE, Bluemke DA, Burke GL, Detrano R, Diez Roux AV, Folsom AR, Greenland P, Jacob DR, Jr., Kronmal R, Liu K, Nelson JC, O’Leary D, Saad MF, Shea S, Szklo M, Tracy RP. Multi-ethnic study of atherosclerosis: Objectives and design. Am J Epidemiol. 2002;156:871–881

15. Ong KL, Morris MJ, McClelland RL, Hughes TM, Maniam J, Fitzpatrick AL, Martin SS, Luchsinger JA, Rapp SR, Hayden KM, Sandfort V, Allison MA, Rye KA. Relationship of lipids and lipid-lowering medications with cognitive function: The multi-ethnic study of atherosclerosis. American journal of epidemiology. 2018;187:767–776

16. Geovanini GR, Wang R, Weng J, Tracy R, Jenny NS, Goldberger AL, Costa MD, Liu Y, Libby P, Redline S. Elevations in neutrophils with obstructive sleep apnea: The multi-ethnic study of atherosclerosis (mesa). International Journal of Cardiology. 2018;257:318–323

17. Fitzpatrick AL, Rapp SR, Luchsinger J, Hill-Briggs F, Alonso A, Gottesman R, Lee H, Carnethon M, Liu K, Williams K, Sharrett AR, Frazier-Wood A, Lyketsos C, Seeman T. Sociodemographic correlates of cognition in the multi-ethnic study of atherosclerosis (mesa). The American journal of geriatric psychiatry: official journal of the American Association for Geriatric Psychiatry. 2015;23:684–697

18. Genuth S, Alberti KG, Bennett P, Buse J, Defronzo R, Kahn R, Kitzmiller J, Knowler WC, Lebovitz H, Lernmark A, Nathan D, Palmer J, Rizza R, Saudek C, Shaw J, Steffes M, Stern M, Tuomilehto J, Zimmet P. Follow-up report on the diagnosis of diabetes mellitus. Diabetes care. 2003;26:3160–3167

19. Levey AS, Stevens LA, Schmid CH, Zhang YL, Castro AF, 3rd, Feldman HI, Kusek JW, Eggers P, Van Lente F, Greene T, Coresh J, Ckd EPI. A new equation to estimate glomerular filtration rate. Ann Intern Med. 2009;150:604–612

20. Hui TH, McClelland RL, Allison MA, Rodriguez CJ, Kronmal RA, Heckbert SR, Michos ED, Barter PJ, Rye KA, Ong KL. The relationship of circulating fibroblast growth factor 21 levels with incident atrial fibrillation: The multi-ethnic study of atherosclerosis. Atherosclerosis. 2018;269:86–91

21. Silverman MG, Harkness JR, Blankstein R, Budoff MJ, Agatston AS, Carr JJ, Lima JA, Blumenthal RS, Nasir K, Blaha MJ. Baseline subclinical atherosclerosis burden and distribution are associated with frequency and mode of future coronary revascularization: Multi-ethnic study of atherosclerosis. JACC Cardiovasc Imaging. 2014;7:476–486

22. Chobanian AV, Bakris GL, Black HR, Cushman WC, Green LA, Izzo JL, Jr., Jones DW, Materson BJ, Oparil S, Wright JT, Jr., Roccella EJ. The seventh report of the joint national committee on prevention, detection, evaluation, and treatment of high blood pressure: The jnc 7 report. Jama. 2003;289:2560–2572

23. Swirski FK, Libby P, Aikawa E, Alcaide P, Luscinskas FW, Weissleder R, Pittet MJ. Ly-6chi monocytes dominate hypercholesterolemia-associated monocytosis and give rise to macrophages in atheromata. The Journal of Clinical Investigation. 2007;117:195–205

24. van der Valk FM, Kuijk C, Verweij SL, Stiekema LCA, Kaiser Y, Zeerleder S, Nahrendorf M, Voermans C, Stroes ESG. Increased haematopoietic activity in patients with atherosclerosis. European Heart Journal. 2017;38:425–432

25. van Leeuwen M, Gijbels MJJ, Duijvestijn A, Smook M, van de Gaar MJ, Heeringa P, de Winther MPJ, Tervaert JWC. Accumulation of myeloperoxidase-positive neutrophils in atherosclerotic lesions in ldlr-/-mice. Arteriosclerosis, Thrombosis, and Vascular Biology. 2008;28:84–89

26. Rotzius P, Thams S, Soehnlein O, Kenne E, Tseng CN, Bjorkstrom NK, Malmberg KJ, Lindbom L, Eriksson EE. Distinct infiltration of neutrophils in lesion shoulders in apoe-/-mice. The American journal of pathology. 2010;177:493–500

27. Doring Y, Soehnlein O, Drechsler M, Shagdarsuren E, Chaudhari SM, Meiler S, Hartwig H, Hristov M, Koenen RR, Hieronymus T, Zenke M, Weber C, Zernecke A. Hematopoietic interferon regulatory factor 8-deficiency accelerates atherosclerosis in mice. Arterioscler Thromb Vasc Biol. 2012;32:1613–1623

28. Drechsler M, Megens RT, van Zandvoort M, Weber C, Soehnlein O. Hyperlipidemia-triggered neutrophilia promotes early atherosclerosis. Circulation. 2010;122:1837–1845

29. Huang Z-S, Chien K-L, Yang C-Y, Tsai K-S, Wang C-H. Peripheral differential leukocyte counts in humans vary with hyperlipidemia, smoking, and body mass index. Lipids. 2001;36:237–245

30. Sarwar N, Danesh J, Eiriksdottir G, Sigurdsson G, Wareham N, Bingham S, Boekholdt SM, Khaw KT, Gudnason V. Triglycerides and the risk of coronary heart disease: 10,158 incident cases among 262,525 participants in 29 western prospective studies. Circulation. 2007;115:450–458

31. Varbo A, Benn M, Tybjaerg-Hansen A, Nordestgaard BG. Elevated remnant cholesterol causes both low-grade inflammation and ischemic heart disease, whereas elevated low-density lipoprotein cholesterol causes ischemic heart disease without inflammation. Circulation. 2013;128:1298–1309

32. Saja Maha F, Baudino L, Jackson William D, Cook H T, Malik Talat H, Fossati-Jimack L, Ruseva M, Pickering Matthew C, Woollard Kevin J, Botto M. Triglyceride-rich lipoproteins modulate the distribution and extravasation of ly6c/gr1(low) monocytes. Cell Reports. 2015;12:1802–1815

33. Smith JD. Dysfunctional hdl as a diagnostic and therapeutic target. Arterioscler Thromb Vasc Biol. 2010;30:151–155

34. Van Lenten BJ, Hama SY, de Beer FC, Stafforini DM, McIntyre TM, Prescott SM, La Du BN, Fogelman AM, Navab M. Anti-inflammatory hdl becomes pro-inflammatory during the acute phase response. Loss of protective effect of hdl against ldl oxidation in aortic wall cell cocultures. J Clin Invest. 1995;96:2758–2767

